# Systems analysis of ribosomal CAR-site dynamics

**DOI:** 10.64898/2026.03.28.714829

**Authors:** Luis Perez, Marc Iradukunda, Daniel Krizanc, Kelly M. Thayer, Michael P. Weir

## Abstract

Developing approaches to link structure and function is an ongoing challenge in computational and structural biology. Using a systems-level framework, we present here an analysis pipeline in a Python package, mdsa-tools, that constructs network representations of structures in a time series of trajectory frames from molecular dynamics (MD) simulations. Here, we demonstrate its use on a ribosomal subsystem. The subsystem is centered on the CAR interaction surface, a “brake pad” adjacent to the aminoacyl (A-site) decoding center that tunes protein translation rates. We leverage unsupervised learning algorithms to explore the conformational landscape of behaviors visited by two versions of the subsystem (brake-on and brake-off) that differ at the codon 3’ adjacent to the A-site codon. Our network representations of MD frames embody H-bond interactions between all pairwise combinations of residues in the system. By utilizing per-frame vector representations of network edges, we can apply standard clustering and dimensionality reduction methods to explore behavioral differences between the brake-on and brake-off versions of the system. K-means clustering of frame vectors revealed a striking separation of the two system versions, consistent with principal components analysis (PCA) embeddings and Uniform Manifold Approximation and Projection (UMAP) embeddings. Dissection of K-means centroids and PCA loadings highlighted H-bond interactions between residue pairs in the ribosome’s peptidyl site (P site), suggesting potential allosteric signaling across the subsystem.

**Author summary:** With the impressive development of computational algorithms to successfully simulate the dynamics of biological molecules over time, the exploration and incorporation of systems modes of analysis is a natural next step to begin to understand the molecular dynamics behaviors that emerge from these experiments. Following the approaches of classical molecular genetics, we used a “computational genetics” paradigm where we introduced changes (mutations) in potentially important residues, changing their identities or modifying their chemical properties, and asked how the dynamic system responded to these changes, viewing the simulations as a series of movie frames of the dynamic structure over time. Starting with network representations of each frame’s structure, where the nodes are residues, and the edges denote H-bond interactions between the residues, we used several unsupervised machine learning algorithms to uncover behavioral changes in the different mutated versions of the system. Applied to our ribosome neighborhood, this revealed unexpected changes in behavior at the ribosome peptidyl site (P site) in response to mutating mRNA residues on the other side of the aminoacyl site (A site) codon, suggesting long-range allosteric interactions across the neighborhood.

## 1. Introduction

Biochemical functions of proteins depend on their structures, which are intimately linked to their amino acid sequences [1]. This relationship also applies to nucleic acid structures and nucleic acid/protein complexes whose conformations are sensitive to H-bond and pi-stacking interactions [2]. Conformational transitions in biological macromolecules are also pivotal in assessing molecular functions, including allosteric regulation, substrate binding, and product release, expanding our understanding of the dynamic relationships between structure and function [3].

With recent advances in computational power, molecular dynamics (MD) simulations [4] are becoming an increasingly successful way to study the conformational dynamics of biomolecular structures in atomic detail. The power of MD allows us to explore the relationships between structure and function and to capture the conformational transitions that may represent behaviors characteristic of different versions of the system of interest. Many methods have been proposed for extracting functionally relevant motions from MD simulations, including clustering approaches paired with appropriate fine-grained metrics [5] and other analytical methods that focus on collective coordinates, low-dimensional variables, or combinations of atomic motions that capture the dominant, functionally relevant conformational changes in the biomolecular structure [3]. Recent development of methods in our group has focused on expanding our “lenses” for exploring systems views of molecular dynamics trajectories, with an emphasis on extracting conformational and behavioral dynamics from MD trajectories. Applying these methods to different versions of a system with altered or substituted residues, in what we think of as a “computational genetics” approach, allows us to pinpoint structural components crucial for functional and conformational behaviors.

Our group has previously employed various measurements, including nucleotide base pi-stacking and backbone atom Cartesian coordinates, as fine-grained metrics from which representations of molecular structures as networks can be derived. These representations can then be leveraged to compare conformational dynamics between different versions of biomolecular systems [6, 7]. In this work, we aim to expand on our previous approaches, with a particular emphasis on H-bond frame-network vector representations. Leveraging these H-bond networks, we apply clustering and dimensionality reduction algorithms to uncover unique conformational behaviors encapsulated in the frames of our MD simulations. Since the frame vector features are mapped to spatially localized properties of the system, dissection of our results provides an opportunity to uncover key structural components associated with these behaviors of the MD system. Using our analysis pipeline and H-bond metric, we uncover potential allosteric changes at the ribosome P-site, as well as throughout the ribosomal subsystem originating from a two-nucleotide mutation at the +1 codon (3’ codon adjacent to the A-site mRNA).

## 2. Results and discussion

### 2.1 An MD subsystem for systems-based analysis

We examined two versions of a ribosome subsystem with localized structural mutations, which we expected could have broad neighborhood-wide effects on behavior, possibly reflecting different functional states and conformations. Our goal was to harness systems framework methodologies and network representations as tools for developing filters (lenses) that highlight differences in conformational behaviors and then localize key parts of the structures underlying these behaviors.

Our 494-residue subsystem of ribosome (Fig 1A) is well suited for this general goal and is based on the 5JUP structure [10] of the translocation stage II yeast ribosome coupled with Taura syndrome virus IRES (internal ribosome entry site), which mimics the mRNA and tRNA. The subsystem is centered around the ribosome’s three-residue CAR surface [7], which behaves as an extension of the A-site tRNA anticodon, acting as a “brake pad” that engages the (+1) codon immediately 3’ of the decoding center A-site codon. We showed previously that the CAR brake pad has two distinctive behaviors, brake-on and brake-off, depending on the sequence at the +1 codon [8, 11, 12]. G-leading +1 codons (+1GNN where N is any nucleotide) have high levels of H-bonding with the CAR surface (brake-on), whereas non-G-leading +1 codons have much lower H-bonding levels (brake-off; Fig 1C), leading to faster protein translation, as supported by analysis of ribosome profiling data [8, 13]. For our analysis, we used two nearly identical subsystems that differed only at the +1 codon, +1GCU (brake-on) and +1CGU (brake-off). Then, to explore the repertoire of behaviors of each subsystem version, we assembled 30 independently initialized replicate trajectories.

**Fig 1.**
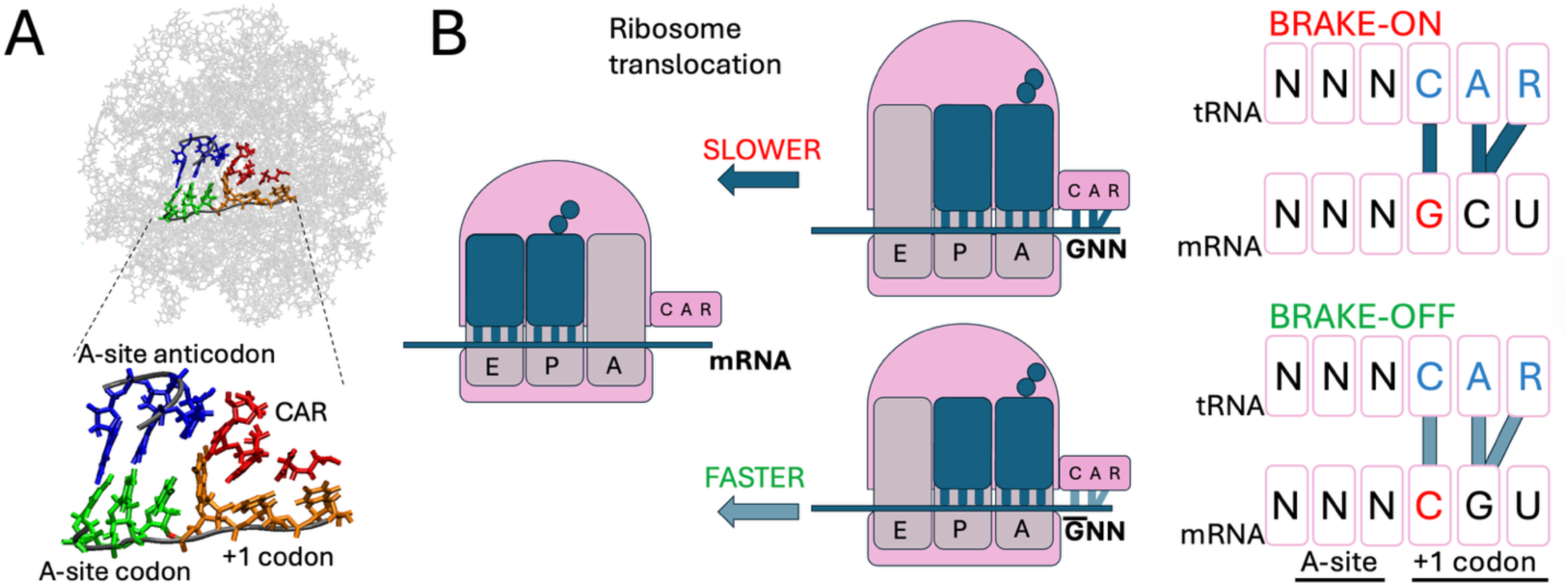
The CAR braking system. (A) We performed molecular dynamics (MD) simulations of a 494-residue subsystem of the yeast ribosome at translocation stage II, translating Taura syndrome virus mRNA [7–9]. We focus on 12 residues at the A and CAR sites. (B) The 3-residue CAR surface is hypothesized to tune translation rates [8]. CAR is conserved and consists of yeast *(S. cerevisiae)* 18S rRNA C1274, A1427 (*E. coli* 16S rRNA C1054, A1196), and yeast Rps3 R146. (C) G-leading +1 codons (brake-on) have higher H-bonding with CAR than +1 non-G-leading codons (brake-off).

### 2.2 A workflow to process network representations of MD trajectory frames

We used network representations of MD trajectory frames to capture broad neighborhood behaviors across the system. Nodes were defined at the level of system residues, although finer-grain levels of resolution, such as at the atomic level, could have been chosen, but with the penalty of significantly increasing the network complexity and corresponding data memory required for each MD frame. The relationships (network edges) between residue nodes could have been defined by different modes of interaction, or combinations of these relationship types, such as H-bond, electrostatic, van der Waals, or pi-stacking interactions. Because the positioning of our system’s atoms and residues is significantly influenced by H-bond interactions between residues, this edge measurement was chosen for our current study.

Each MD frame was represented as a vector in which entries (features) correspond to the H-bond count for a specific residue pair (see Section 3.2). This is equivalent to vectorizing (flattening) the upper triangle of the residue-residue interaction matrix into a single per-frame vector representation. The per-frame H-bond counts (per residue pair) were computed using MDTraj [14], a Python library for analysis of MD trajectories, by assessing each MD frame’s atomic coordinates for the presence of a given H-bond. An H-bond was identified for a pair of residues when they were sufficiently close to each other and with appropriate geometry to interact through H-bonding (θ>120°; H-acceptor distance < 2.5 Å; see Section 3.2).

Our ordered frame vectors together define a matrix (Fig 2), representing a data encapsulation of MD behavior through a lens (metric) of choice. By representing the MD frames as vectors, we opened the opportunity to apply vector-based analyses to compare and dissect the key properties of the information embedded in each frame. Because the 1050 frame-vector features (after dropping empty columns; see Section 3.2) map to spatially localized properties of the system, analysis of the frame vectors provided an opportunity to uncover key structural components associated with important behaviors of the MD system. The workflow (Fig 2) offers multiple modes of analysis focusing on conformational behavior assessment through clustering and low-dimensional embeddings that uncover MD frame relationships and key structural properties related to structure-function assessments.

**Fig 2.**
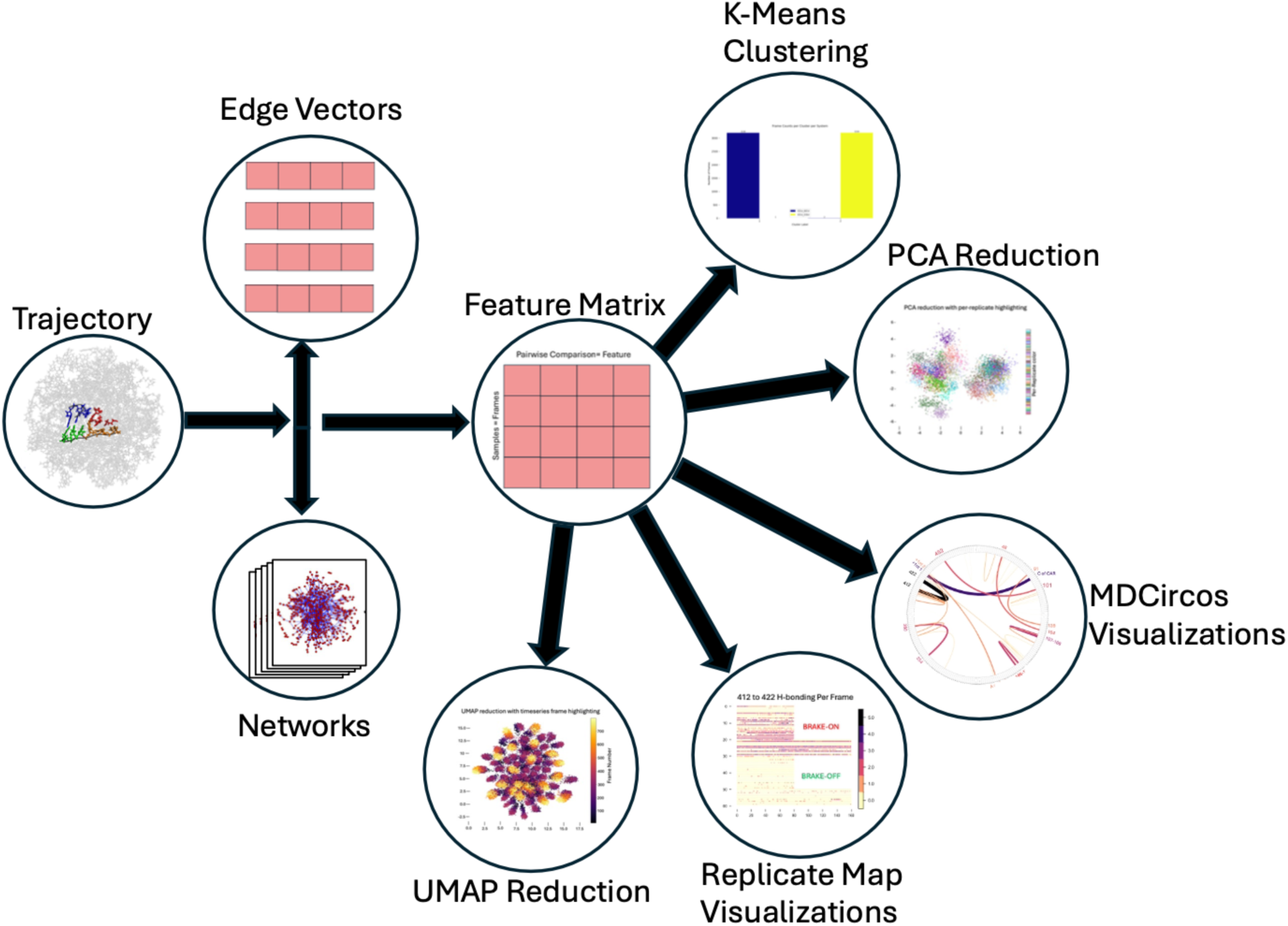
MD trajectory analysis pipeline. From left to right, we begin with a trajectory file and convert it into a set of networks (one for each trajectory frame) that could be represented as either networks or edge vectors. These vectors are vertically concatenated to create a matrix that can then be used as input to unsupervised machine-learning algorithms such as K-means, PCA, or UMAP. Results can be visualized using scatter plots, MDCircos plots, or MD replicate maps (Fig 5).

### 2.3 K-means clustering of MD frame vectors

The K-means clustering algorithm is highly effective in determining the optimal cluster assignments for a given value of k (the number of clusters) and has been used successfully to group MD frames with similar characteristics. The frames, input as vectors, are commonly clustered based on the Cartesian coordinates of selected backbone atoms. However, other representations of frames, such as vectorized residue-residue contact maps with varying metrics, including pi-stacking and electrostatic interactions, can also be utilized [5, 6, 15, 16].

We applied the K-means algorithm to the frame vectors of H-bond data, where the vectors for both versions of the ribosome subsystem (brake-on and brake-off) were combined into a single set of vectors that the algorithm would cluster without knowledge of the subsystem version origin of each vector. After running the algorithm with multiple choices of k (the number of clusters), silhouette analysis (Fig 3B), which measures within-cluster cohesion (the average distance of each data point to other points in its cluster; here, each data point is a frame vector) and separation (average distance to vectors in the nearest other cluster), revealed k=2 to be an optimal choice of k. Strikingly, with k=2, rather than a “salt and pepper” result, the frame vectors were cleanly separated into two clusters based on their subsystem version of origin (Fig 3A).

**Fig 3.**
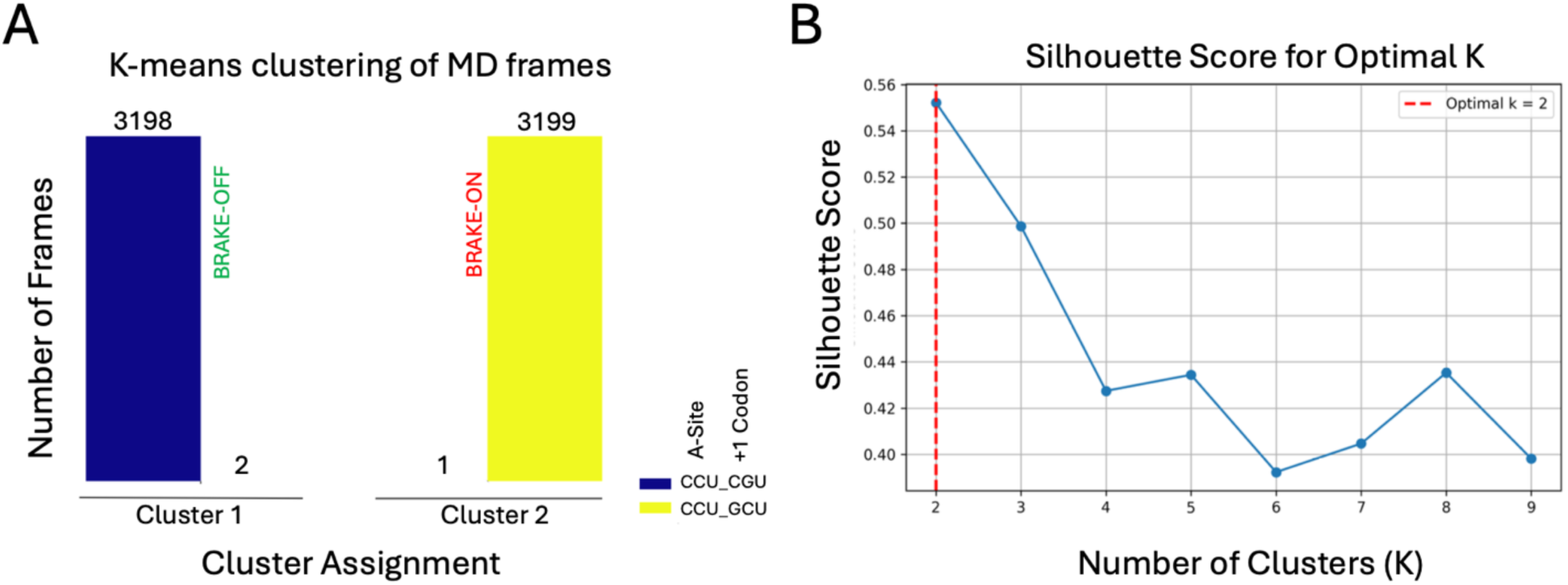
K-means clustering of MD frames separates the frames of the brake-on and brake-off structures. (A) K-means clustering was performed on the combined set of brake-on and brake-off frame vectors. Cluster assignments are denoted below each set of bars, brake-off frame identities are highlighted in blue and brake-on in yellow, with counts above each bar. (B) Silhouette score analysis supports exploring k=2 as the optimal number of clusters (k).

To interpret this striking separation, we used the final centroids computed by the K-means algorithm (see Section 3.3). Each centroid represents the average of all frames in that cluster (here, a component-wise mean vector), and the resulting clusters in our analysis corresponded nearly entirely to a specific system (brake-on vs brake-off; Fig 3). Thus, it follows naturally that taking the difference between the two centroids and inspecting which features had the largest differences would identify residue pairs that tend to be different between the two system versions (S1 Table; Section 3.3 for details). Notably, the 3 highest absolute differences pointed at mRNA-tRNA H-bond interactions in the P-site (411-422 and 412-422) as well as between the C of CAR, and the first nucleotide of the +1 codon (94-426), indicating their importance in defining the cluster separation. Interestingly, the 411-422 average count was higher in the brake-on system (further discussed in Section 2.4), and the 412-422 interaction was higher in the brake-off system. This hints that there may be a switch in hydrogen bonding patterns when mutations are made to the incoming codon (3’ adjacent to the A-site).

The separation of our structures into two distinct clusters, using residue H-bond data, parallels our previous applications of K-means clustering to the same MD simulations using atomic Cartesian coordinate data [7] or pi-stacking data [6], where in all three cases, a clean separation of the two versions of the ribosomal subsystem was observed. These equivalent clustering results obtained using different network edge values as residue interaction measures strengthened confidence in the significance of the overall results. It also suggested that it is not crucial to have identified the “best” interaction measurement for edge values in a systems analysis of MD trajectories if the utilized edge values reflect behavior variation across the system.

Indeed, these observations also emphasize that MD trajectories represent a simulation of molecular dynamics where atom behaviors are determined by force fields optimized to reflect actual atomic relationships. The force fields determine atom placements in successive frames, and their placements are then interpreted by assessment algorithms such as hbond in AmberTools’ cpptraj [17] or MDTraj’s baker hubbard function [14] to infer H-bond or other interactions. So effectively, the MD simulations are inferring rather than measuring the interaction strengths, but these inferred measurements provide excellent lenses to assess system behavior.

### 2.4 Dimensional reduction of MD frame vectors

Our implementation of the K-means clustering algorithm partitions the high-dimensional vectors into clusters based on Euclidean distances. Other unsupervised algorithms that reduce the dimensionality of the data can also uncover edge relationships that are particularly influential in determining the system behaviors. Principal component analysis (PCA) and Uniform Manifold Approximation and Projection (UMAP; see Section 2.6) embeddings were used to reduce the data into two dimensions, thereby providing a transformation of the data designed to be easier to visualize (in 2D scatter plots) and that may make differences between the assessed MD systems easier to spot.

Using PCA to analyze the combined sets of brake-on and brake-off MD frames, we projected (Fig 4) our high-dimensional vectors (or feature matrix) with 1,050 features down to a lower-dimensional subspace by extracting the first two principal components (eigenvectors of the covariance matrix of the data), which accounted for 5.4% and 2.1% of the total explained variance respectively, well within the range of typical dynamic coordinate analysis [3]. The same frame vectors were analyzed using UMAP, which provides a non-linear embedding that emphasizes local relationships amongst the projected vectors (Section 2.6). As with K-means clustering, the application of the PCA algorithm resulted in a strikingly clean separation of frame vectors for the brake-on and brake-off versions of the MD system, rather than a salt and pepper mix (Fig 4A).

**Fig 4.**
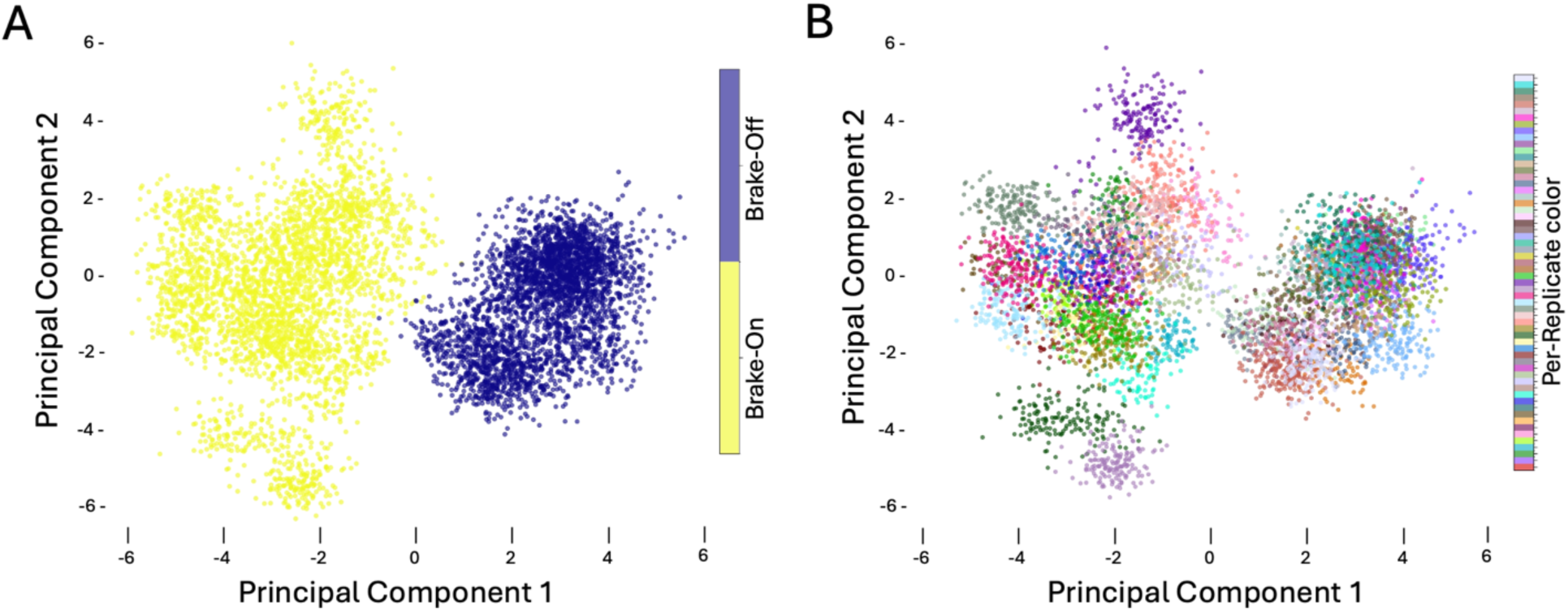
Comparing MD frames. (A) Both the resulting cluster assignments from K-means clustering (Fig 3) and the first principal component of PCA analysis separate the brake-on and brake-off versions of the system, suggesting distinct structural conformations and behaviors. (B) 60 independently initiated MD replicates occupy distinct but overlapping conformational/behavioral spaces in the embedding space created by PCA analysis. Each color pertains to a specific replicate.

### 2.5 Dissection of principal component coefficients reveals influential residue interactions

The embedding of vectors in PC1/PC2 space involves computing weights (eigenvector loadings) for each element of the vectors. Hence, in our analysis, there is a weight associated with each pair of residues for each principal component. The vector feature weights with the highest magnitudes (see methods) are most influential in determining placement along the PC1 (or PC2) axis. Since our two versions of the MD system (brake-on and brake-off) are well separated along the PC1 axis, it follows that the vector features (residue pairs) with the most prominent weights tend to show the largest differences in H-bonding between the brake-on and brake-off behaviors.

Influential vector features were identified by rank ordering the magnitudes of feature loading weights (the coefficients of the eigenvectors; S2-S3 Table) for PC1 of the PCA embedding. The influential vector features were visualized using chord diagrams (MDCircos), where the thicknesses of arcs conveyed the strengths of positive and negative weightings (Fig 5A). Generally, the residue pairs with higher H-bonding in the brake-off MD trajectories had positive weightings, and the pairs with higher H-bonding in the brake-on version of the system had negative weightings. When mapped back to the ribosome subsystem, these residue pairs with the greatest differences between the two versions of our structures were distributed throughout the subsystem (Fig 5A; S1 Fig). Since the two versions of the system only differed by two nucleotides at the +1 codon (3’ adjacent to the A-site codon), the broad distribution of implicated residue pairs suggested allosteric communication across the structure.

**Fig 5.**
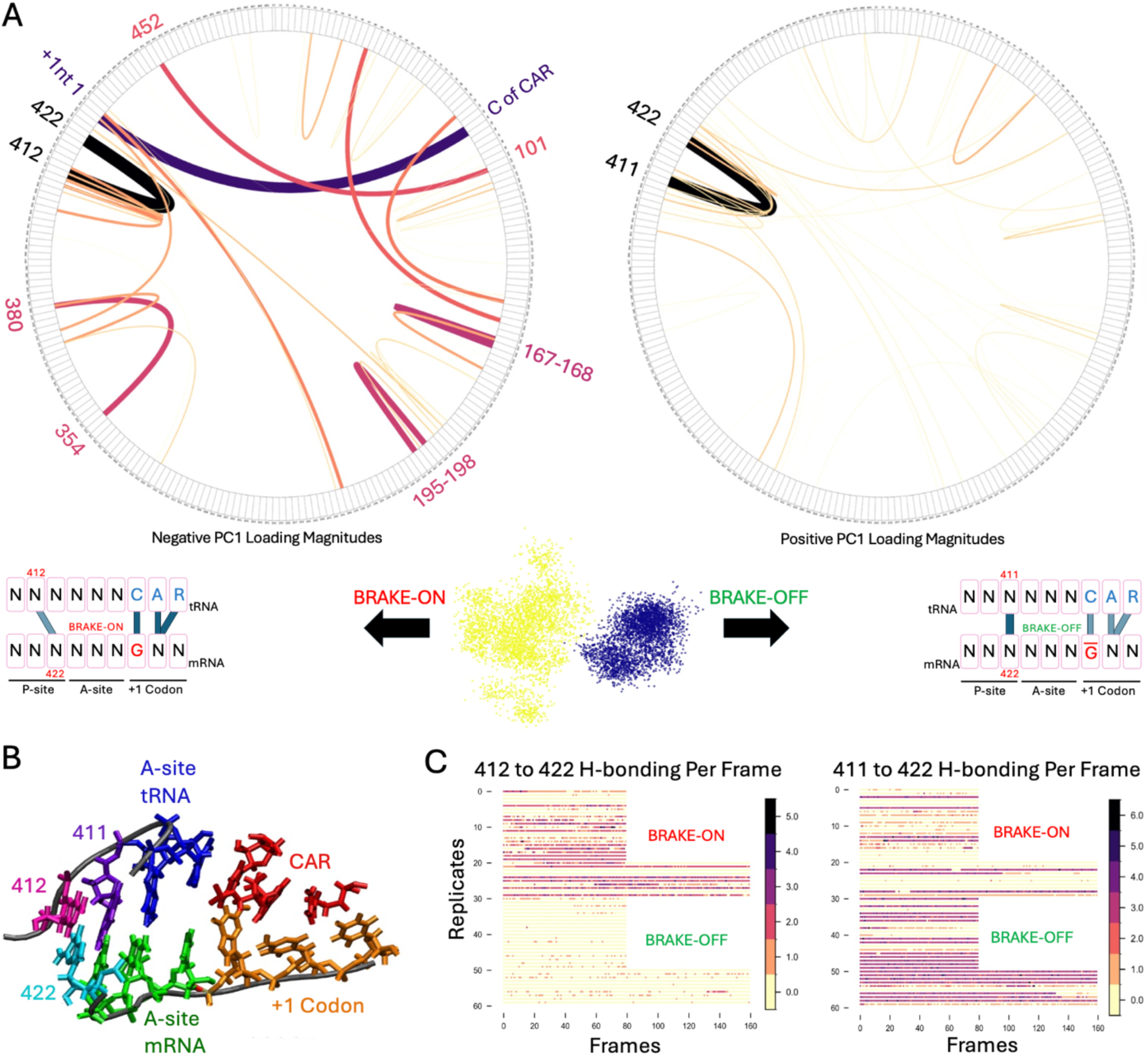
PC1 loadings uncover residue interactions in the P-site. (A) MDCircos plots depict PCA feature weightings that reveal residue pairs with large differences in H-bonding between brake-on and brake-off versions of the system. Thicker, darker lines depict higher magnitude weights. The left-hand circle shows negative weights, and the right-hand circle shows positive weights. Depicted beside each MDCircos plot are schematics illustrating the observed H-bonding strengths by CAR and the P-site residues of interest. (B) IRES P-site residues with strong H-bonding in the system’s brake-on (412-422) or brake-off (411-422) versions. (C) Time series plots of MD replicates showing elevated H-bonding for 411-422 (brake-off) and 412-422 (brake-on). The x-axis shows frames 1-160 corresponding to 20-100 ns of trajectories (2 frames/ns).

The residue pairs with the highest-magnitude PC1 loadings were 411-422 and 412-422 (Fig 5A, 5B; S2 Table). Notably, these were the same residue pairs highlighted by dissecting our K-means centroids (see Section 2.3), unexpectedly mapping to the peptidyl (P) site of the ribosome subsystem and reinforcing our results. When the CAR brake is off, there is standard base-pair H-bonding between the P-site wobble (3^rd^) nucleotide of the codon (422) with its corresponding nucleotide in the tRNA anticodon (411, tRNA nt34). However, when the CAR brake is on, the wobble nucleotide (422) instead base pairs more strongly with the adjacent (out-of-phase) nucleotide of the anticodon (412; tRNA nt35). Because our subsystem includes a viral internal ribosome entry site (IRES) RNA that mimics mRNA and tRNA (see methods), this behavior warrants future investigation in structures with authentic tRNA anticodons.

To further assess the behavior difference at the ribosome’s P site, we used a Replicate Map representation (Fig 5C) to examine the 411-422 and 412-422 H-bonding behaviors over time across each replicate MD trajectory. This revealed that the brake-off version of the subsystem rarely exhibits the out-of-phase (412-422) base pairing, whereas most of the brake-on replicates show this base pairing at least some of the time. In-phase base pairing (411-422) is consistently observed in the brake-off replicates but is infrequent in most of the brake-on replicates. Notably, the out-of-phase base pairing is typically weaker (fewer H-bond counts between atoms) in the brake-off version of the system than in the brake-on version of the system.

### 2.6 UMAP embeddings reveal time-dependent MD paths

Algorithms for projecting high-dimensional data into two dimensions use computations focusing on different aspects of the relationships between the embedded vectors [18]. The powerful UMAP algorithm [19] emphasizes details of local relationships between a vector and its immediate neighbors, in contrast to the PCA algorithm, which emphasizes global relationships. We were interested in how the local relationships exposed by the UMAP algorithm would display the frame vectors of our brake-on and brake-off MD trajectories.

Two crucial hyperparameters for the UMAP algorithm are the number of neighbors and the minimum distance. The first parameter balances the preservation of local structure and global structure by varying the number of points being used to compute the local neighborhood (see methods). The second parameter, minimum distance, controls how tightly UMAP packs points together in the low-dimensional embedding; smaller values pertain to tighter bunches and larger values scatter vectors. In practice, the minimum distance parameter mainly controls how tightly points are packed in the two-dimensional embedding, whereas the number of neighbors more strongly influences how the relationships between vector points are embedded.

Strikingly, UMAP projections of the brake-on and brake-off frame vectors showed a clean separation of the brake-on and brake-off versions of the MD system (Fig 6A) across a range of parameters, supporting the utility of UMAP in addition to PCA embeddings to assess MD behaviors. Based on hyperparameter sweeps (S2 Fig), we choose reductions with the neighbor hyperparameter set to 915 and minimum distance set to 1.0 to balance local and global relationships. The grouping of replicate-specific frames was not surprising since the close-by adjacent frames in replicates were emphasized by the UMAP algorithm. In contrast, the PCA embeddings showed that the replicates tended to occupy localized but partially overlapping clouds in the embedding space (Fig 4B).

**Fig 6.**
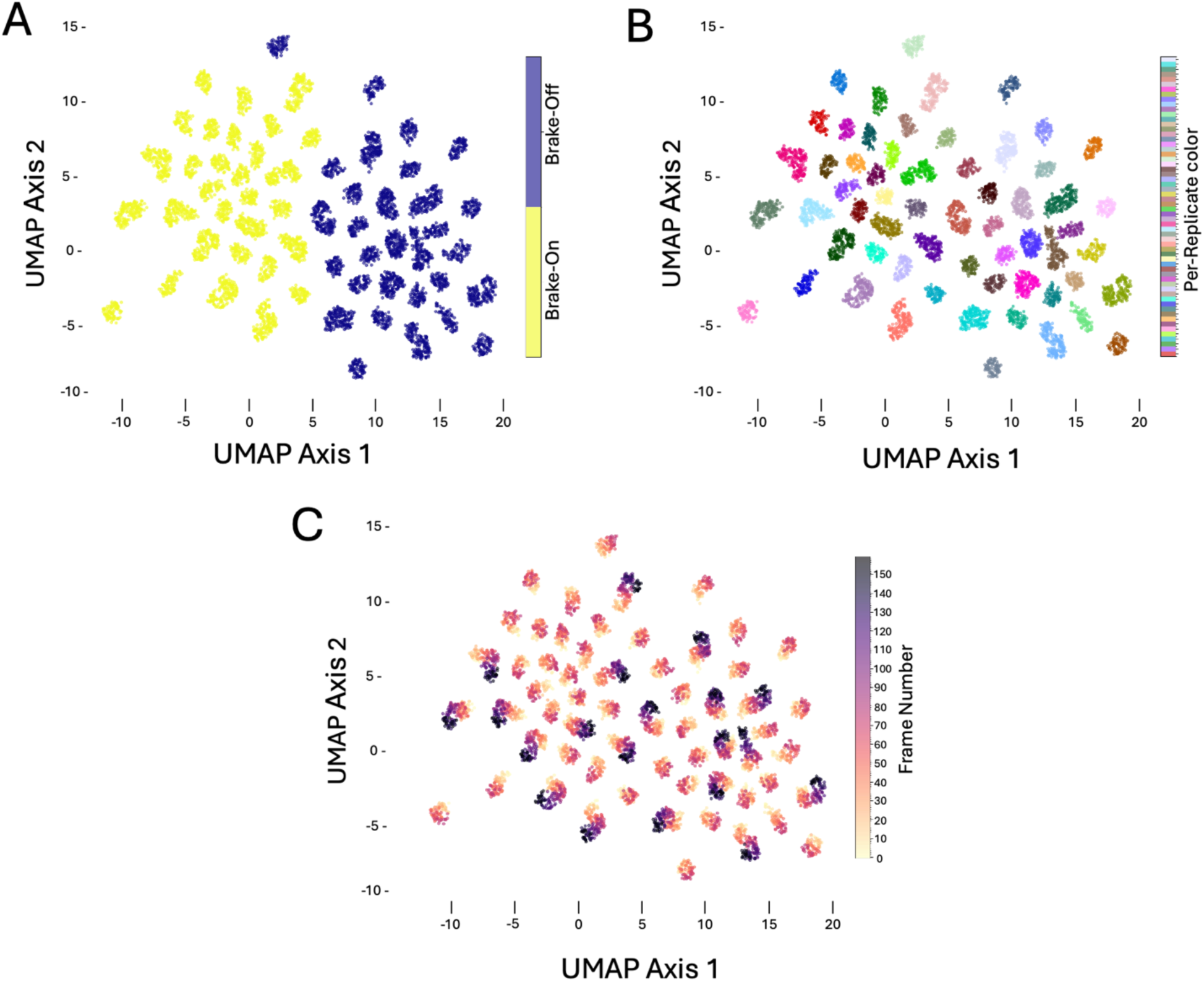
UMAP embeddings reveal time-dependent MD paths. (A) UMAP embedding highlighting two systems: brake-on (yellow) and brake-off (blue). (B) UMAP embedding of brake-on and brake-off versions of the MD system with independently initiated replicates highlighted as in Fig 4B. (C) UMAP embedding of brake-on and brake-off versions of the MD system with trajectory frames heat-mapped sequentially. The later frames of the longer (100ns) replicates are deep purple.

Interestingly, when embeddings were run with all production frames (without stripping the first 20ns; see Section 3.1; S3 Fig), a stronger temporal trend is visible. In these embeddings, the replicates are also divided across the first UMAP axis (Fig 6A; S3A Fig), with brake-on replicates typically having UMAP axis 1 coordinate values greater than ∼6.0 and brake-off having UMAP axis 1 coordinate values less than ∼6.0. Per-replicate timeseries heatmapping of trajectory frames (S3B-C Fig) shows many of the early frames from replicates of both versions of the system are in a more central position along the UMAP 1 axis (closer to 5.0) and then extend further into the system-specific region of the embedding (brake-on towards 15.0, brake-off towards -2.5). Earlier frames in the temporal paths of each replicate often overlap, suggesting that some of the replicates initially visit similar conformations.

Both our UMAP and PCA embeddings support the value of performing multiple independent MD simulations (in our case, 30 replicates for each system version) to improve sampling of the conformational space visited by each system. By using frames from multiple replicate trajectories for each version of our system, we built a fuller picture of the embedding spaces visited by each version, which are strikingly separated for the two versions.

### 2.7 Conclusions

Consistent with the hypothesis that the two versions of our system may have different structural conformations [6, 7], the frames of each of our structures were strikingly separated by K-means, PCA, and UMAP. UMAP’s temporal view in particular (Fig 6; S3 Fig) revealed that the replicates of the brake-on and brake-off versions of the system evolve towards separate regions of the embedding space, supporting the hypothesis that they may be settling into distinct structural conformations. Both our K-means and PCA analyses (supported by temporal assessment of H-bond patterns; Fig 5C) revealed that the most pronounced H-bond pattern differences were at the P-site as well as at other neighborhood locations far from the mutated codon at the mRNA entry site (S1 Fig). This suggests the presence of allosteric interactions across the dynamic structure and is of interest for future study by the group. As mentioned, our model uses a viral IRES RNA to mimic the mRNA-tRNA context, and validation with authentic ribosome complexes will be needed to confirm these allosteric patterns [10].

Additionally, using the workflow provided by our Python package mdsa-tools, we have shown that dynamic molecular behaviors from MD simulations can be effectively analyzed when each frame is stored as a vector of network edge values. In general, it seems that different lenses (pi-stacking, atomic coordinates, H-bond counts) represent the biology well enough to distinguish the two states, brake-on and brake-off, but specific interaction metrics may affect which localized differences are pinpointed [6, 7]. While we have used pairwise H-bond counts between residues as edge values, this work builds on the use of other edge measures to describe molecular systems in the literature [5, 6, 15], and the assessment of weighted combinations of these measures may be valuable for future studies.

## 3. Methods

### 3.1 Molecular Dynamics (MD) simulations

MD simulations of our subsystem of the ribosome were performed as described previously [7]. Briefly, we simulated a 494-residue subsystem of the ribosome based on a cryo-EM structure of the ribosome at translocation stage II (PDB ID: 5JUP)[9, 10]. Shown in Fig 1A, our structure includes a viral internal ribosome entry site (IRES) RNA that mimics mRNA and tRNAs in our systems. The N2 neighborhood consists of a 35 Å-radius sphere centered on C1054 (the C of CAR). The outermost residues are restrained during dynamics with a harmonic potential force 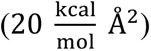 to maintain the structural conformation of translocation stage II.

For each +1 codon identity modification of interest, nucleotide identities were changed using AMBER22’s tLEaP [17], and 30 independently initiated replicate simulations (20x60ns; 10x100ns; sampled at 2 frames per ns) were performed. We discarded the first 20ns of each replicate as RMSD assessments of RNA and protein backbone atoms [7] indicated the system had settled by this time. For this analysis, we concatenated all replicates of each version of the system, producing system version-specific concatenated trajectories encompassing a broad range of conformational behaviors.

The results in this study involve two N2 neighborhood variants that illustrate the CAR interaction surfaces’ varying behavior based on the identity of the codon at the +1-codon position, the codon 3’ adjacent to the A-site codon. These structures contained 5’CCU in the A-site mRNA, 5’GGG in the A-site tRNA anticodon, and 5’GCU or 5’CGU in the +1-codon position and were denoted as tGGG_aNNU_+1GCU and tGGG_aNNU_+1CGU, respectively. In this work, we refer to tGGG_aNNU_+1GCU as the “brake-on” and tGGG_aNNU_+1CGU as the “brake-off” versions of our system, reflecting the hypothesized brake mechanism of the CAR interaction surface [8].

### 3.2 Generation of systems representations

The results presented in this work utilize H-bond networks generated with MDTraj [14], and we also provide a module that can take as input matrices generated with the *hbond* command and the *uuseries* flag using AmberTools cpptraj [17]. For the Python-native network generation, the module expects either trajectories and their respective topologies, a pre-loaded MDTraj trajectory object, or a Protein Data Bank (PDB) file as input. The full list of allowed input file formats is available in MDTraj’s documentation and includes DCD (CHARMM/NAMD), XTC/TRR (GROMACS), LAMMPS trajectories, and MDCRD files (AMBER). Once trajectories are loaded, H-bond counts between all atoms present in the trajectory are calculated with the parameters defined in the Baker-Hubbard method [14, 20] (angle θ > 120°, distance between the hydrogen atom and the acceptor atom < 2.5 Å). This processing step pools H-bonds found between any two atoms on different residues and creates an adjacency matrix with row and column indices equivalent to the residue indices defined in the topologies of the trajectories. The edge weight between two residues is the total count of H-bonds found between them (regardless of which atoms on either residue were involved in the interaction). Each adjacency matrix is then flattened in row-major order to create a vector representation of each frame, keeping only the upper triangle and discarding redundant symmetric data. Finally, we vertically concatenate the vector representations of each frame to create a matrix where each frame (row) is represented as a vector of its given H-bond edge values, and each column is a specific, distinct pair of residues. As a pre-processing step, we removed all zero columns. This reduced the dimensionality of our data from 26,106 columns to 1,050 columns. This filtering did not alter the relative geometry of the data in feature space for any distance-based or linear method but improved memory and general runtime efficiencies.

In the cpptraj-based workflow, the input is a matrix where each row corresponds to a particular frame of a trajectory, and each column denotes an H-bond interaction between two atoms in that frame. The atoms represented in each column are dependent on the mask the user provides in the original run. For consistency, each row is rebuilt by aggregating all atom-atom interactions between a given pair of residues into a single “H-bond” count. Because heavy computations are done by the well-optimized AMBER library for H-bond identification, this workflow is recommended for dense sampling or long timescale trajectories. A noteworthy difference is that the H-bond identification parameters are different from those of the Baker Hubbard method (angle θ >135°; distance between the donor atom and the acceptor atom < 3.0 Å), but were of no consequence for the results presented here, for which we used the Baker Hubbard definition, but provide the CPPTRAJ branch as an alternative for future workflows.

The result from either workflow is a set of frames encapsulated as network edge-weight vectors. As discussed above, we used the brake-on and brake-off (+1 codon G-leading and non-G-leading) versions of the N2 neighborhood as benchmarks. These vectors can be stored as NumPy arrays [21], or NumPy’s zip archive arrays, which allow for easy manipulation and integration into other scientific computing packages. Importantly, the final matrix used in this analysis contains all vectors from each system version so that downstream unsupervised learning analyses operate on the combined set of vectors and can more readily identify structural differences between the two system versions.

Although in this study we focused on AMBER trajectories of a ribosomal subsystem, the mdsa-tools package also includes fixtures as part of our continuous integration (CI) tests that use other MD engines and systems [22, 23], demonstrating that the pipeline is not specific to the ribosome case study. Our toolkit builds on various examples in the literature of similar MD analysis software with different metrics and pipelines [6, 16, 24, 25]

### 3.3 K-means clustering of MD frames

The matrix (described above) is a standard input format for many implementations of classical machine-learning algorithms, including the package used here, scikit-learn [26]. Available within this package, K-means is an iterative clustering algorithm used to partition N objects into K disjoint and non-empty subsets. It achieves this in several steps: first, K data points (vectors) are randomly assigned as initial centroids of K clusters; then each data point is assigned to the cluster with the nearest centroid based on a distance metric; then the centroids are updated to the average of all data points in their cluster; next the previous two steps are repeated for a set number of iterations or until we reach convergence [27]. In this study, we treated each edge vector as a data point and used the Euclidean distance between these vectors when applying the K-means algorithm.

The K-means clustering algorithm expects a parameter, K, the number of clusters. Different approaches can be used to select an appropriate number of clusters, and for this study, we used a silhouette score analysis [28]. Silhouette scores are calculated for individual samples (in our case, frames) using the mean intra-cluster distance (a) and the mean nearest-cluster distance (b) for each data point. The silhouette coefficient for a sample is (b – a)/ max (a, b). Samples have a score closer to +1 when the clusters are compact and well separated, and closer to 0 when they are not. To summarize the data, we report the mean of all the sample silhouette scores to calculate a “mean silhouette score” of all the clustered points. K-means clustering was performed on these vector representations of the data, and the algorithm was run for k = 2 to 10 for silhouette score cross-validation, which suggested an optimal choice of k = 2 (Fig 3B).

While K-means does not report weights in the same way that the PCA algorithm does (See Section 3.4), one can take advantage of the centroids and assigned labels to further interpret the resulting clusters. Because we had only two clusters, and the frames of each system version (brake-on; brake-off) were assigned almost entirely to a unique cluster (Fig 3), interpretation of centroids was akin to interpretation of a single system version. Thus, by taking the difference and absolute difference between each centroid (so component-wise differences) and examining the centroid values feature-by-feature to find the largest differences, we can identify which residue-residue interactions were the most different between the two system versions (S1 Table). Signed difference indicates which cluster has the higher average value for that feature, which is particularly useful because our clusters map directly to the system versions. Here, we subtracted the brake-off dominant centroid from the brake-on dominant centroid, resulting in signed differences where a negative value indicates a higher average H-bond count in brake-off and a positive value indicates a higher average H-bond count in brake-on. We also took the absolute difference, which simply indicates the biggest differences between the two clusters, regardless of which system version has the higher value.

### 3.4 Dimensional reduction of frame vectors

Like our implementation of K-means clustering, we used scikit-learn to run PCA. The goal of PCA is to find axes onto which we can project our data that capture the greatest variance. Briefly, this is achieved by constructing the covariance matrix of our original matrix, which is a symmetric matrix where every entry (i, j) represents the covariance between features (columns) i and j. Then, we compute the eigenpairs (λ_(_, v_(_) of the covariance matrix. Finally, the eigenpairs are sorted by decreasing eigenvalue, and we use the two eigenvectors corresponding to the largest eigenvalues (i.e., the maximum and second largest eigenvalues) as the axes onto which we project our data [29]. These are our “principal components” that maximize variance, as each eigenvalue equals the variance along its eigenvector. For this study, we ran our analyses on the same matrix used for K-means clustering, with the number of principal components set to n = 2, in effect reducing the dimensionality from 1,050 unique columns (features [27]) to 2, while preserving the greatest differences (variance) in our data.

UMAP, a dimensionality reduction technique rooted in manifold learning, is commonly employed in Python through the umap-learn package [19]. First, a K-nearest neighbors’ graph is constructed with nodes as samples in the high-dimensional feature space. The edges between them can be thought of as similarity scores (fuzzy similarities) based on distance (here Euclidean) and scaled by density around each point. Then, UMAP projects the points into a lower-dimensional space using spectral embedding and re-creates the K-nearest neighbors’ graph by progressively updating (optimizing) the positions of the data points to mirror the high-dimensional neighborhood structure. We ran our reductions with the specifications of 915 neighbors and 1.0 minimum distance, which were found to be preferred for balancing local and global relationships (since UMAP’s K-nearest-neighbors approach already emphasizes local relationships; S2 Fig)[30].

One of the benefits of using a linear method like principal component analysis, as opposed to non-linear methods like UMAP, is the ability to easily extract coefficients (loadings) from the eigenvectors of the covariance matrix (the principal-component axes). These coefficients inform us of how strongly each feature contributes to a principal component axis. Recalling the structure of our edge vectors (Fig 2), each feature (column) in our matrix represents H-bond counts between specific pairs of residues; therefore, features (residue pairs) with more prominent loadings (see below) contribute the most to variance in the new principal component axes.

The most pronounced separation of our two system versions is across the midpoint (0) of the first principal component (Fig 4A), so it follows that, if we examine the weightings (or contributions) of each feature in defining these axes, their sign (-/+) will give us an indication of which direction (left/right) along the first principal component they are contributing to placement of frames, and their magnitude (weighting^’^) reflects the relative importance in determining this placement along the given principal component axis. Our embedding space visualizations are simple scatter plots generated with matplotlib [31], where we provide the coordinates of each frame in the embedding space defined by the two new principal components. As described above, for each system version, we analyzed 3200 frames of MD simulation data, which were plotted in the principal components space as scatterplots (Fig 4).

### 3.5 MDCircos plots

MDCircos plots can be thought of as chord diagrams that mirror Circos plots [32] (Fig 5A), a common tool in genomics. The implementation in this study was a natural extension of the workflow, making use of the Python package Pycircos [33]. We begin by taking a Pandas data frame created as output in the analysis module and use it as the basis for the creation of Circos plots. The MDCircos plots take all residue indices found in the data frame and use them as identifiers for sections of the outer circle (called arcs). Then it draws lines between each one of the arcs with color and thickness scaled to the value of the magnitude of the eigenvector coefficients of the principal components. It simultaneously creates two plots, one for the first principal component and one for the second.

For this project, we created two separate MDCircos plots for the first principal component. We created one MDCircos plot for all the negative loadings (squaring for magnitude readings) and separately, one for all the positive loadings. Because the brake-on and brake-off versions of our system were well separated along the first principal component (Fig 4A), the negative PC1 loadings mainly highlight residue pairs with higher H-bond counts in the brake-on version of the system, and the positive PC1 loadings highlight residue pairs with higher H-bond counts in the brake-off version of the system (Fig 5C).

### 3.6 MDReplicate maps

MDReplicate maps provide a useful tool to visualize a measurement of interest across time in independently initialized replicate MD experiments, akin to a line plot but using color instead of height. These plots provide a visual overview of the stabilities and instabilities of measurements, and whether different MD replicates tend to show similar or distinct behaviors. The tool takes as input an array of labels that correspond to each frame (row) in the original matrix. These labels can represent H-bond counts, K-means clustering frame assignments, or even principal component weightings. The second parameter is a list that denotes how frames are distributed amongst their original trajectories (because our matrix contains vectors from all replicates and systems, see Section 3.2).

Using this input, the tool generates NumPy arrays of coordinates and color identifiers and uses these arrays to plot a color-mapped scatterplot using matplotlib. The color represents the value of the associated label, providing a clear visualization of temporal change. For the implementation shown in Fig 5C, we provide a Python list denoting the distribution of frames contributed by each replicate, as well as the corresponding list of H-bond counts between residues of interest.

## Acknowledgements

We thank Joe Coolon, Scott Holmes, Amy MacQueen, Dalton Soper, Mia Wichman, Miki Lynch, Pete Hwang, Eric Sakkas, and Mitsu Raval for engaging in discussions and Henk Meij for technical assistance with high-performance computing.

## Data & code availability

All Analysis code and figure generation scripts are available inside our GitHub Repository (https://github.com/zeper-eng/mdsa-tools) in the manuscript_data_availability subdirectory. Detailed documentation and tutorials for using mdsa-tools as a Python package are also present in the repository. MD restart structures and topology files, as well as code for molecular dynamics experiments, can be found at https://doi.org/10.25438/wes02.25328632.v1 [8]. Additionally, the software environment for the figures and data produced in this work includes Python 3.13.5 using NumPy 2.2.6, pandas 2.3.1, SciPy 1.16.1, scikit-learn 1.7.1, umap-learn 0.5.9.post2, Matplotlib 3.10.5, MDTraj 1.10.3, MDAnalysis 2.9.0, and python-circos 0.3.0. Trajectory preparation/processing was performed with AmberTools22 as described in [8].

## Supporting information captions

S1 Fig. Structural Visualization of the top 20 (Blue) pairwise residue H-bond interactions rank-ordered by loading magnitude from PCA analysis

S2 Fig. Hyperparameter Sweeps (Grid-Search) for “optimal” UMAP parameters.

S3 Fig. UMAP embeddings of replicates without stripping early frames typically excluded from analysis

S1 Table. Top 20 absolute differences between the two centroids computed by K-means clustering.

S2 Table. Top 20 Principal Components Weightings (rank ordered by PC1 magnitude)

S3 Table. Top 20 Principal Components Weightings (rank ordered by PC2 magnitude)

## References

1. Alberts B, Johnson A, Lewis J, Raff M, Roberts K, Walter P. Molecular biology of the cell. 4th ed. New York: Garland Science; 2002.

2. Bottaro S, Di Palma F, Bussi G. The role of nucleobase interactions in RNA structure and dynamics. Nucleic Acids Res. 2014;42(21):13306–13314. doi: 10.1093/nar/gku972.

3. Amadei A, Linssen AB, Berendsen HJ. Essential dynamics of proteins. Proteins. 1993;17(4):412–425. doi: 10.1002/prot.340170408.

4. Hollingsworth SA, Dror RO. Molecular dynamics simulation for all. Neuron. 2018;99(6):1129–1143. doi: 10.1016/j.neuron.2018.08.011.

5. Thayer KM, Lakhani B, Beveridge DL. Molecular dynamics–Markov state model of protein ligand binding and allostery in CRIB-PDZ: Conformational selection and induced fit. J Phys Chem B. 2017;121(22):5509–5514. doi: 10.1021/acs.jpcb.7b02083.

6. Sakkas ED, Krizanc D, Thayer KM, Weir MP. StACKER, a tool for systems level analysis of base stacking in nucleotide-rich structures. BioRxiv [Preprint]. 2025 bioRxiv 648419 [posted 2025 Apr 13; cited 2025 Nov 4]. Available from: https://www.biorxiv.org/content/10.1101/2025.04.13.648419v1 doi: 10.1101/2025.04.13.648419.

7. Dalgarno C, Scopino K, Raval M, Nachmanoff C, Sakkas ED, Krizanc D, et al. The CAR–mRNA interaction surface is a zipper extension of the ribosome A site. Int J Mol Sci. 2022;23(3):1417. doi: 10.3390/ijms23031417.

8. Sun J, Hwang P, Sakkas ED, Zhou Y, Perez L, Dave I, et al. GNN codon adjacency tunes protein translation. Int J Mol Sci. 2024;25(11):5914. doi: 10.3390/ijms25115914.

9. Scopino K, Dalgarno C, Nachmanoff C, Krizanc D, Thayer KM, Weir MP. Arginine methylation regulates ribosome CAR function. Int J Mol Sci. 2021;22(3):1335. doi: 10.3390/ijms22031335.

10. Abeyrathne PD, Koh CS, Grant T, Grigorieff N, Korostelev AA. Ensemble cryo-EM uncovers inchworm-like translocation of a viral IRES through the ribosome. eLife. 2016;5:e14874. doi: 10.7554/eLife.14874.

11. Scopino K, Williams E, Elsayed A, Barr WA, Krizanc D, Thayer KM, et al. A ribosome interaction surface sensitive to mRNA GCN periodicity. Biomolecules. 2020;10(6):849. doi: 10.3390/biom10060849.

12. Barr WA, Sheth RB, Kwon J, Cho J, Glickman JW, Hart F, et al. GCN sensitive protein translation in yeast. PLoS One. 2020;15(9):e0233197. doi: 10.1371/journal.pone.0233197.

13. Wu CC, Zinshteyn B, Wehner KA, Green R. High-resolution ribosome profiling defines discrete ribosome elongation states and translational regulation during cellular stress. Mol Cell. 2019;73(5):959–970. doi: 10.1016/j.molcel.2018.12.009.

14. McGibbon RT, Beauchamp KA, Harrigan MP, Klein C, Swails JM, Hernández CX, et al. MDTraj: A modern open library for the analysis of molecular dynamics trajectories. Biophys J. 2015;109(8):1528–1532. doi: 10.1016/j.bpj.2015.08.015.

15. Serçinoğlu O, Ozbek P. gRINN: A tool for calculation of residue interaction energies and protein energy network analysis of molecular dynamics simulations. Nucleic Acids Res. 2018;46(W1):W554–W562. doi: 10.1093/nar/gky381.

16. Franke L, Peter C. Clustering and analyzing ensembles of residue interaction networks from molecular dynamics simulations. J Chem Inf Model. 2025;65(20):11203–11214. doi: 10.1021/acs.jcim.5c01298.

17. Case DA, Aktulga HM, Belfon KAA, Ben-Shalom IY, Berryman JT, Brozell SR, et al. Amber 2022 reference manual [Internet]. San Francisco (CA): University of California, San Francisco; 2022. [cited 2025 Nov 4]. Available from: https://ambermd.org/doc12/Amber22.pdf.

18. Trozzi F, Wang X, Tao P. UMAP as a dimensionality reduction tool for molecular dynamics simulations of biomacromolecules: A comparison study. J Phys Chem B. 2021;125(19):5022–5034. doi: 10.1021/acs.jpcb.1c02081.

19. McInnes L, Healy J, Saul N, Großberger L. UMAP: uniform manifold approximation and projection. J Open Source Softw. 2018;3(29):861. doi: 10.21105/joss.00861.

20. Baker EN, Hubbard RE. Hydrogen bonding in globular proteins. Prog Biophys Mol Biol. 1984;44(2):97–179. doi: 10.1016/0079-6107(84)90007-5.

21. Harris CR, Millman KJ, van der Walt SJ, Gommers R, Virtanen P, Cournapeau D, et al. Array programming with NumPy. Nature. 2020;585(7825):357–362. doi: 10.1038/s41586-020-2649-2.

22. Seyler S, Beckstein O. Molecular dynamics trajectory for benchmarking MDAnalysis; 2017 [cited 2025 Dec 31]. Database: figshare [Internet]. Available from: 10.6084/m9.figshare.5108170.v1.

23. Beckstein O, Gowers R, Alibay I, Fan S, Wang L, Matta M. MDAnalysis/MDAnalysisData: 0.9.0 (release-0.9.0) [Internet]. Zenodo; 2023 [cited 2025 Dec 31]. Available from: 10.5281/zenodo.10058664.

24. Pérez-Hernández G, Hildebrand PW. Mdciao: Accessible analysis and visualization of molecular dynamics simulation data. PLoS Comput Biol. 2025;21(4):e1012837. doi: 10.1371/journal.pcbi.1012837.

25. Kožić M, Bertoša B. Trajectory maps: Molecular dynamics visualization and analysis. NAR Genom Bioinform. 2024;6(1):lqad114. doi: 10.1093/nargab/lqad114.

26. Pedregosa F, Varoquaux G, Gramfort A, Michel V, Thirion B, Grisel O, et al. Scikit-learn: Machine learning in Python. J Mach Learn Res. 2011;12:2825–2830.

27. Bishop CM. Pattern recognition and machine learning. New York (NY): Springer; 2006.

28. Rousseeuw PJ. Silhouettes: A graphical aid to the interpretation and validation of cluster analysis. J Comput Appl Math. 1987;20:53–65. doi: 10.1016/0377-0427(87)90125-7.

29. Abdi H, Williams LJ. Principal component analysis. Wiley Interdiscip Rev Comput Stat. 2010;2(4):433–459. doi: 10.1002/wics.101.

30. Venna J, Kaski S. Neighborhood preservation in nonlinear projection methods: An experimental study. In: Dorffner G, Bischof H, Hornik K, editors. Artificial Neural Networks—ICANN 2001. Berlin, Heidelberg: Springer; 2001. pp. 485–491. doi: 10.1007/3-540-44668-0_68.

31. Hunter JD. Matplotlib: A 2D graphics environment. Comput Sci Eng. 2007;9(3):90–95. doi: 10.1109/MCSE.2007.55.

32. Krzywinski M, Schein J, Birol İ, Connors J, Gascoyne R, Horsman D, et al. Circos: An information aesthetic for comparative genomics. Genome Res. 2009;19(9):1639–1645. doi: 10.1101/gr.092759.109.

33. ponnhide. pyCircos: Circos plot in matplotlib. Version 0.3.0 [Internet]. San Francisco: GitHub; 2022 [cited 2025 Nov 4]. Available from: https://github.com/ponnhide/pyCircos.

