## Supplemental Figures and Tables for "Systems analysis of ribosomal CAR-site dynamics"

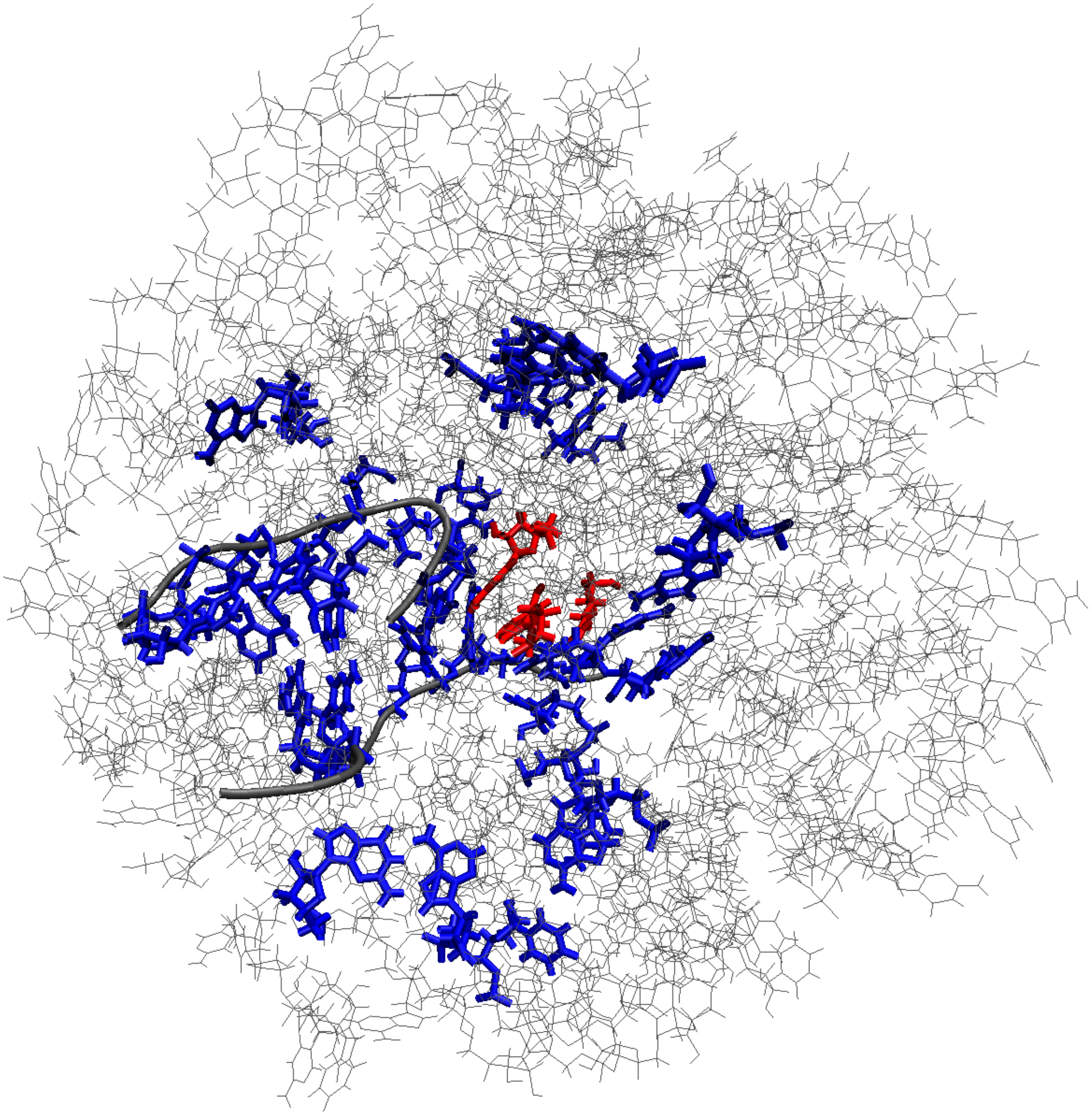

**S1 Fig.** Structural Visualization of the top 20 (Blue) pairwise residue H-bond interactions rank-ordered by loading magnitude from PCA analysis. The CAR residues are visualized in red, the backbone of the viral IRES that mimics mRNA and tRNA is shown in tube form, and all other unrestrained residues of the structure are represented with thin lines.

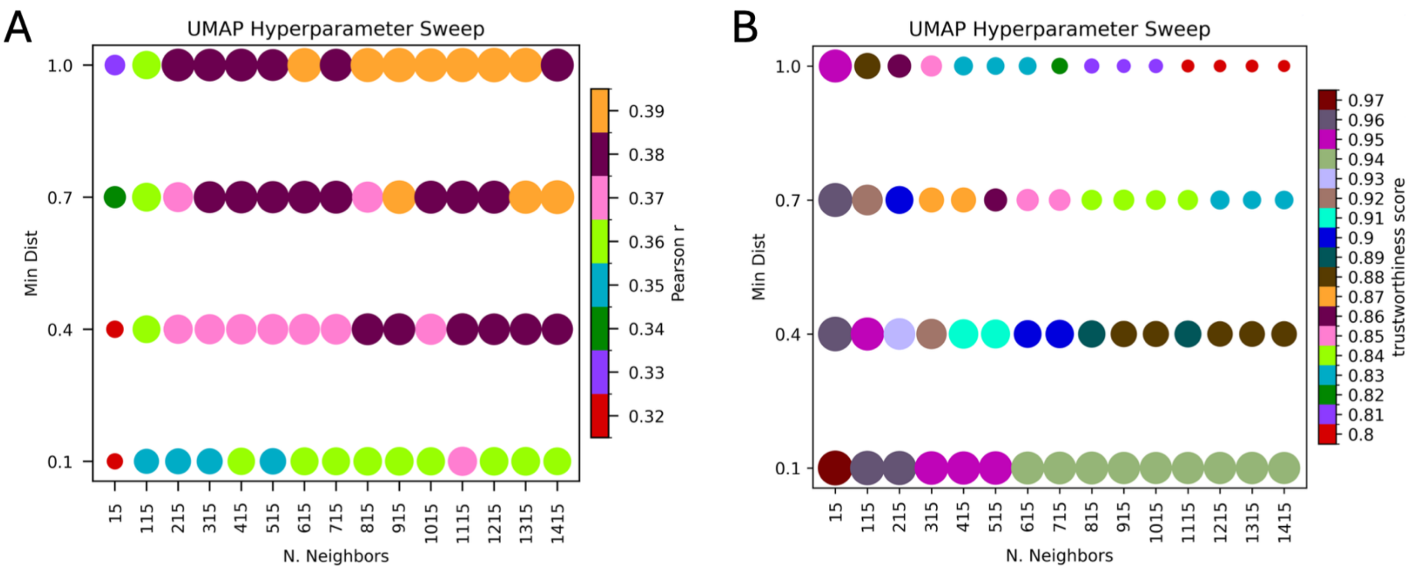


**S2 Fig.** Hyperparameter Sweeps (Grid-Search) for “optimal” UMAP parameters. (A) We scanned varying neighborhood sizes from 15-1415 and .1 minimum distance to 1.0 minimum distance using a Pearson correlation coefficient to compare pairwise distances (Euclidean) between our high-dimensional frame vectors and equivalent pairwise distances of our low-dimensional (UMAP reduced) frame vectors to assess global neighborhood preservation. (B) For local neighborhood optimization, the metric trustworthiness [30] was used to compare how well the k-nearest neighbors relationships in our original feature space are preserved in the low-dimensional embedding (k=15; here number of neighbors being evaluated). Trustworthiness penalizes points that appear among the k nearest neighbors in the embedding but were not neighbors in the original feature space, with penalties being proportional to how far in rank the points were outside the neighborhood. (A, B) All results were plotted in bubble plot form with bubble color and size scaling with their respective Pearson correlation coefficient or trustworthiness values. Based on these results, for our main analysis, we set N = 915 neighbors and minimum distance = 1.0 to balance preservation of local (B) and global (A) relationships.

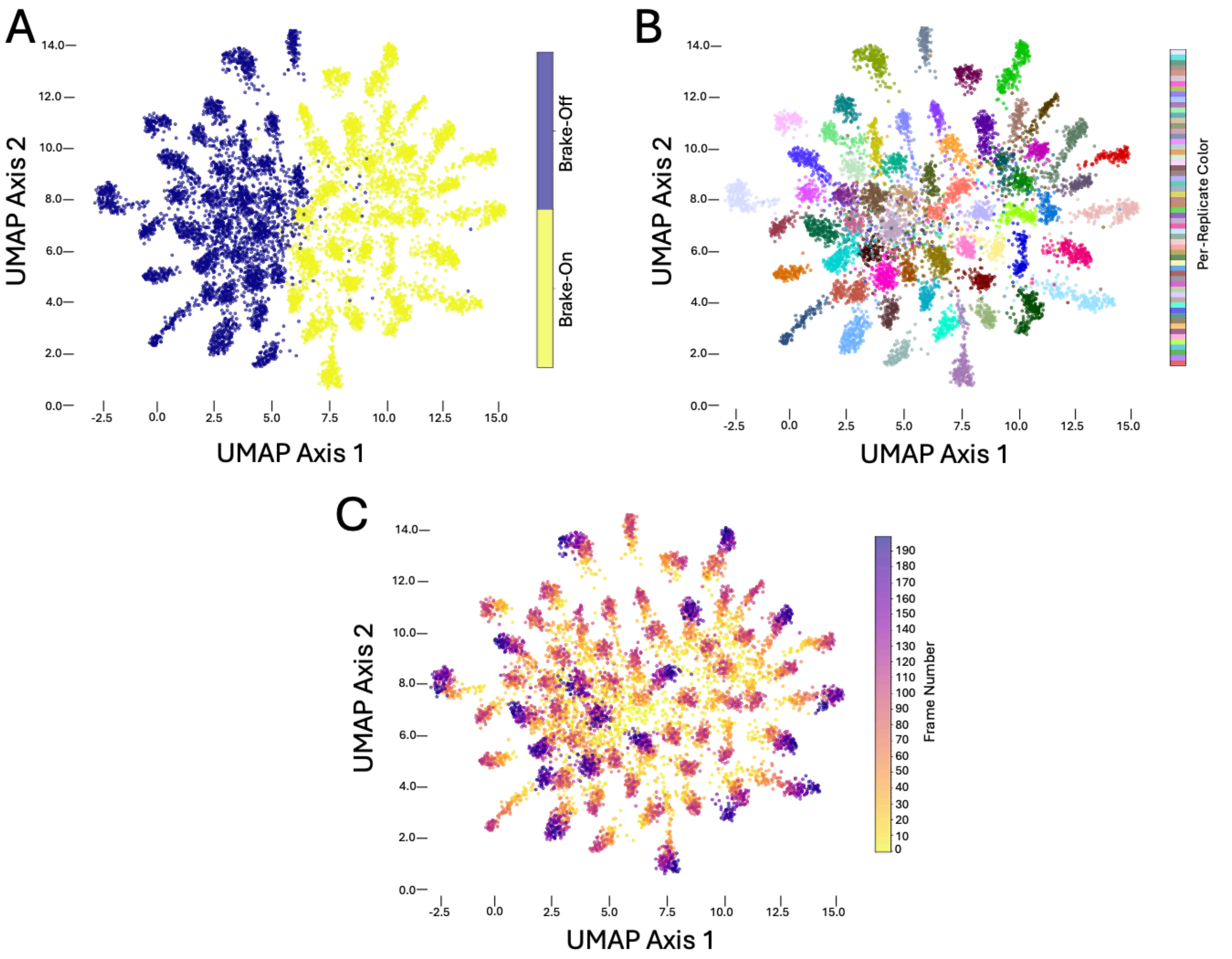


**S3 Fig.** UMAP embeddings of replicates without stripping early frames typically excluded from analysis (see Section 3.1). (A) UMAP embedding highlighting two systems: brake-on (yellow) and brake-off (blue). (B) UMAP embedding of brake-on and brake-off versions of the MD system with independently initiated replicates highlighted as in Fig 4B. (C) UMAP embedding of brake-on and brake-off versions of the MD system with trajectory frames heat-mapped sequentially. The later frames of the longer (100ns) replicates are deep purple.

S1 Table. Top 20 absolute differences between the two centroids computed by K-means clustering.

| Residue pairs | Difference* | Absolute Difference |
|---|---|---|
| 411-422 | -1.224 | 1.224 |
| 94-426 | 1.063 | 1.063 |
| 412-422 | 1.024 | 1.024 |
| 195-198 | 0.802 | 0.802 |
| 354-380 | 0.78 | 0.78 |
| 101-452 | 0.772 | 0.772 |
| 167-168 | 0.722 | 0.722 |
| 48-164 | 0.632 | 0.632 |
| 91-139 | 0.619 | 0.619 |
| 376-426 | 0.618 | 0.618 |
| 240-427 | 0.608 | 0.608 |
| 408-425 | -0.606 | 0.606 |
| 125-126 | -0.594 | 0.594 |
| 412-413 | -0.594 | 0.594 |
| 422-423 | -0.592 | 0.592 |
| 328-403 | -0.572 | 0.572 |
| 94-425 | -0.57 | 0.57 |
| 66-75 | -0.57 | 0.57 |
| 26-467 | -0.566 | 0.566 |
| 166-167 | -0.545 | 0.545 |

* Difference in average H-bonding (brake-on - brake-off)

S2 Table. Top 20 Principal Components Weightings (rank ordered by PC1 magnitude)

| Comparisons | PC1 Loadings | PC1 Magnitude* |
|---|---|---|
| 411-422 | 0.333 | 0.111 |
| 412-422 | -0.213 | 0.045 |
| 94-426 | -0.191 | 0.037 |
| 167-168 | -0.151 | 0.023 |
| 195-198 | -0.146 | 0.021 |
| 354-380 | -0.143 | 0.021 |
| 422-423 | 0.133 | 0.018 |
| 101-452 | -0.135 | 0.018 |
| 66-75 | 0.133 | 0.018 |
| 48-164 | -0.129 | 0.017 |
| 408-425 | 0.121 | 0.015 |
| 91-139 | -0.118 | 0.014 |
| 240-427 | -0.114 | 0.013 |
| 328-403 | 0.113 | 0.013 |
| 412-413 | 0.113 | 0.013 |
| 410-411 | -0.112 | 0.012 |
| 125-126 | 0.105 | 0.011 |
| 406-413 | -0.104 | 0.011 |
| 374-380 | -0.101 | 0.01 |
| 376-426 | -0.099 | 0.01 |

* Loading squared

S3 Table. Top 20 Principal Components Weightings (rank ordered by PC2 magnitude)

| Residue pairs | PC2 Loadings | PC2 Magnitude* |
|---|---|---|
| 411-422 | 0.444 | 0.197 |
| 409-424 | -0.222 | 0.049 |
| 66-75 | 0.201 | 0.04 |
| 167-168 | 0.198 | 0.039 |
| 376-426 | 0.182 | 0.033 |
| 406-413 | 0.149 | 0.022 |
| 410-411 | -0.145 | 0.021 |
| 425-426 | 0.142 | 0.02 |
| 328-403 | 0.137 | 0.019 |
| 424-425 | 0.129 | 0.017 |
| 324-379 | 0.127 | 0.016 |
| 166-169 | -0.126 | 0.016 |
| 96-240 | -0.121 | 0.015 |
| 124-488 | -0.123 | 0.015 |
| 191-423 | -0.116 | 0.013 |
| 384-388 | -0.113 | 0.013 |
| 240-427 | 0.111 | 0.012 |
| 379-424 | -0.104 | 0.011 |
| 234-242 | -0.1 | 0.01 |
| 190-191 | 0.099 | 0.01 |

*Loading squared
